# Phylogenetic Analysis of SARS-CoV-2 Genomes in Turkey

**DOI:** 10.1101/2020.05.15.095794

**Authors:** Ogün Adebalı, Aylin Bırcan, Defne Çırcı, Burak İşlek, Zeynep Kilinç, Berkay Selçuk, Berk Turhan

## Abstract

COVID-19 has effectively spread worldwide. As of May 2020, Turkey is among the top ten countries with the most cases. A comprehensive genomic characterization of the virus isolates in Turkey is yet to be carried out. Here, we built a phylogenetic tree with globally obtained 15,277 severe acute respiratory syndrome coronavirus 2 (SARS-CoV-2) genomes. We identified the subtypes based on the phylogenetic clustering in comparison with the previously annotated classifications. We performed a phylogenetic analysis of the first thirty SARS-CoV-2 genomes isolated and sequenced in Turkey. We suggest that the first introduction of the virus to the country is earlier than the first reported case of infection. Virus genomes isolated from Turkey are dispersed among most types in the phylogenetic tree. We find two of the seventeen sub-clusters enriched with the isolates of Turkey, which likely have spread expansively in the country. Finally, we traced virus genomes based on their phylogenetic placements. This analysis suggested multiple independent international introductions of the virus and revealed a hub for the inland transmission. We released a web application to track the global and interprovincial virus spread of the isolates from Turkey in comparison to thousands of genomes worldwide.

## 1. Introduction

Severe acute respiratory syndrome coronavirus 2 (SARS-CoV-2) has emerged in Wuhan (Li, et al. 2020), spread across continents and eventually resulted in the COVID-19 pandemic. Although there are significant differences between the current and previously known SARS-CoV genomes, the reason behind it’s pandemic behaviour is still unclear. Genome sequences around the world were revealed and deposited into public databases such as GISAID (Shu and McCauley 2017). With those genomic datasets, it is possible, in fact crucial to reveal the evolutionary events of SARS-CoV-2 to understand the types of the circulating genomes as well as in which parts of the genome differ across these types.

The SARS-CoV-2 virus is homologous to SARS-CoV, and its closer versions were characterized in bats and pangolins (Li, et al. 2020). The virus has been under a strong purifying selection (Li, et al. 2020). With the isolates obtained so far, the sequences of SARS-CoV-2 genomes showed more than 99.9% percent identity indicating a recent shift to the human species (Tang, et al. 2020). Yet, there are unambiguous evolutionary clusters in the genome pool. Various studies use SNP (Tang, et al. 2020) or entropy (Zhao, et al. 2020) based methods to identify evolving virus types to reveal genomic regions responsible for transmission and evolution. Tang et. al identified S and L types among 103 SARS-CoV-2 genomes based on two SNPs at ORF1ab and ORF8 regions which encode replicase/transcriptase and ATF6, respectively (Tang, et al. 2020). The entropy-based approach generated informative subtype markers from 17 informative positions to cluster evolving virus genomes (Zhao, et al. 2020). Another study defined a competitive subtype based on the D614G mutation in the spike protein which facilitates binding to ACE2 to receptor on the host cell surface (Bhattacharyya, et al. 2020). Although whether there is any effect of D614G substitution on the transmissibility is inconclusive (van Dorp, et al. 2020), this mutation has been one of the landmarks for major groupings of the virus family.

In this work, we used publicly available SARS-CoV-2 genome datasets. We aligned the sequences of more than 15,000 whole genomes and built a phylogenetic tree with the maximum likelihood method. We clustered the genomes based on their clade distribution in the phylogenetic tree, identified their genomic characteristics and linked them with the previous studies. We further analysed clusters, mutations and transmission patterns of the genomes from Turkey.

## 2. Materials and methods

To perform our analyses we retrieved virus genomes, aligned them to each other and revealed the evolutionary relationships between them through phylogenetic trees. We assigned the clusters based on the mutations for each genome. We further analyzed the phylogenetic tree with respect to neighbor samples of our genomes of interest to identify possible transmission patterns.

### 2.1. Data retrieval, multiple sequence alignment and phylogenomic tree generation

The entire SARS-CoV-2 genome sequences, along with their metadata were retrieved from the GISAID database **(Table-S1)** (Shu and McCauley 2017). We retrieved the initial batch of genomes (3,228) from GISAID on 02/04/2020. We used Augur toolkit to align whole genome sequences using mafft algorithm (--reorder --anysymbol – nomemsave) (Katoh and Standley 2016). The SARS-CoV2 isolate Wuhan-Hu-1 genome (GenBank:NC_045512.2) was used as a reference genome to trim the sequence and remove insertions in the genomes. Since the initial batch, the new sequences in GISAID were periodically added to the pre-existing multiple sequence alignment (--existing-alignment). The final multiple sequence alignment (MSA) contained 15,501 genomes that were available on May 1^st^ 2020. In the metadata file, some genomes lacked month and day information and contained the year of the sample collection date. The genomes with incomplete metadata were filtered out and the unfiltered MSA consisted of 15,277 sequences. Maximum likelihood phylogenetic tree was built with IQ-TREE with the following options: -nt AUTO (on a 112-core server) -m GTR -fast. Augur was used to estimate the molecular clock through TimeTree (Sagulenko, et al. 2018). For the sample EPI-ISL-428718 we additionally built a separate maximum likelihood phylogenetic tree by using IQ-TREE multicore version 1.6.1 with Ultra-fast Bootstrapping option and 1000 bootstraps.

The sub-tree consisting of Turkey isolates **(Table-1)** were retrieved from the master time-resolved tree by removing the rest of the genomes with the ‘Pruning’ method from ete3 toolkit (Huerta-Cepas, et al. 2016). The tree was visualized in FigTree v1.4.4 (http://tree.bio.ed.ac.uk/software/figtree/), and rerooted by selecting EPI_ISL_428718 as an outgroup. The branch lengths of EPI-ISL-417413 and EPI-ISL-428713 samples were shortened for better visualization. Ggtree (Yu, et al. 2017) package in R was used to generate the tree and corresponding clusters.

**Table 1.** The genome sequences identified in Turkey. See the Supplementary Table – S1 for the full list. All authors are listed in the acknowledgments in detail. The genomes are sorted by the sample collection date.

| Accession | Date | City | Lab | Authors |
| --- | --- | --- | --- | --- |
| EPI_ISL_429866 | 3/16/20 | Afyon | Ministry of Health Turkey | Bayrakdar et al. |
| EPI_ISL_417413 | 3/17/20 | Ankara | Ministry of Health Turkey | Bayrakdar et al. |
| EPI_ISL_424366 | 3/17/20 | Kayseri | Erciyes University | Pavel et al. |
| EPI_ISL_428712 | 3/17/20 | Karaman | Ministry of Health Turkey | Bayrakdar et al. |
| EPI_ISL_429867 | 3/17/20 | Balikesir | Ministry of Health Turkey | Bayrakdar et al. |
| EPI_ISL_429868 | 3/17/20 | Eskisehir | Ministry of Health Turkey | Bayrakdar et al. |
| EPI_ISL_429869 | 3/17/20 | Konya | Ministry of Health Turkey | Bayrakdar et al. |
| EPI_ISL_428716 | 3/18/20 | Ankara | Ministry of Health Turkey | Bayrakdar et al. |
| EPI_ISL_428713 | 3/18/20 | Ankara | Ministry of Health Turkey | Bayrakdar et al. |
| EPI_ISL_428715 | 3/18/20 | Nevşehir | Ministry of Health Turkey | Bayrakdar et al. |
| EPI_ISL_428714 | 3/18/20 | Kastamonu | Ministry of Health Turkey | Bayrakdar et al. |
| EPI_ISL_429865 | 3/18/20 | Çanakkale | Ministry of Health Turkey | Bayrakdar et al. |
| EPI_ISL_428717 | 3/19/20 | Kocaeli | Ministry of Health Turkey | Bayrakdar et al. |
| EPI_ISL_428718 | 3/19/20 | Kocaeli | Ministry of Health Turkey | Bayrakdar et al. |
| EPI_ISL_428719 | 3/21/20 | Siirt | Ministry of Health Turkey | Bayrakdar et al. |
| EPI_ISL_428720 | 3/21/20 | Ankara | Ministry of Health Turkey | Bayrakdar et al. |
| EPI_ISL_428721 | 3/21/20 | Ankara | Ministry of Health Turkey | Bayrakdar et al. |
| EPI_ISL_428722 | 3/22/20 | Balıkesir | Ministry of Health Turkey | Bayrakdar et al. |
| EPI_ISL_428723 | 3/22/20 | Aksaray | Ministry of Health Turkey | Bayrakdar et al. |
| EPI_ISL_429870 | 3/22/20 | Sakarya | Ministry of Health Turkey | Bayrakdar et al. |
| EPI_ISL_429861 | 3/22/20 | Ankara | Ministry of Health Turkey | Bayrakdar et al. |
| EPI_ISL_429862 | 3/22/20 | Ankara | Ministry of Health Turkey | Bayrakdar et al. |
| EPI_ISL_429863 | 3/22/20 | Sakarya | Ministry of Health Turkey | Bayrakdar et al. |
| EPI_ISL_429864 | 3/22/20 | Sakarya | Ministry of Health Turkey | Bayrakdar et al. |
| EPI_ISL_429871 | 3/23/20 | Ankara | Ministry of Health Turkey | Bayrakdar et al. |
| EPI_ISL_429873 | 3/23/20 | Kocaeli | Ministry of Health Turkey | Bayrakdar et al. |
| EPI_ISL_429872 | 3/25/20 | Kocaeli | Ministry of Health Turkey | Bayrakdar et al. |
| EPI_ISL_427391 | 4/13/20 | İstanbul | GLAB | Karacan et al. |
| EPI_ISL_428368 | 4/16/20 | İstanbul | GLAB | Karacan et al. |
| EPI_ISL_428346 | 4/17/20 | İstanbul | GLAB | Karacan et al. |

### 2.2. Genome clustering

We generated phylo-clusters with TreeCluster (Balaban, et al. 2019) which is specifically designed to group viral genomes. The tool supports different clustering options and we used the default option, Max Clade, which identifies clusters based on two parameters, “-t” and “-s”. These parameters define the threshold that two leaf nodes can be distant from each other and assign a minimum support value that connects two leaf nodes or clades, respectively. For this analysis, we only used the distance threshold. The Max Clade algorithm requires leaves to form a clade and satisfy the distance threshold. The number of clusters that can be generated using a phylogenetic tree depends on the pairwise leaf distance cutoff. We manually searched for a meaningful cutoff for the number of phylo-clusters and phylo-subgroups based on their similarity with the previously reported clusters (see below). We used the -t parameter as 0.0084 and 0.00463 for phylo-clusters and phylo-subclusters, respectively. After retrieving the groupings from TreeCluster, we eliminated clusters containing less than 100 sequences (except one sub-cluster with 99 sequences). We categorized those clusters having less than 100 sequences as not clustered. As a result, we obtained four primary and seventeen sub-clusters.

L/S types of the SARS-CoV-2 genomes were previously defined based on the nucleotides at 8782^nd^ and 28144^th^ positions (Tang, et al. 2020). We categorized “TC” and “CT” haplotypes S and L type, respectively. In the cases both these positions correspond to a gap, the sequences were classified N type. All other cases were categorized as unknown types. 614 G/D clustering was applied based on the amino acid at the 614^th^ position of the Spike protein (Jaimes, et al. 2020). Combinations of the nucleotides at positions 241;1059; 3037; 8782; 11083; 14408; 14805; 17747; 17858; 18060; 23403; 25563; 26144; 28144; 28881; 28882; 28883 determined the subtypes for barcode clustering. Sequences that belong to the ten major subtypes (with more than 100 sequences) which constitute 86 percent of all sequences were labelled with their respective 17 nucleotides (Zhao, et al. 2020). All other sequences were categorized as unknown for barcode classification. Six major clusters (Morais Júnior, et al. 2020) were assigned by the previously determined twelve positions (3037; 8782; 11083; 14408; 17747; 17858; 18060; 23403; 28144; 28881; 28882; 28883). The lineages were assigned using the proposed nomenclature by Rabaut et al. through Pangolin COVID-19 Lineage Assigner web server (Rambaut, et al. 2020). Sequences that cannot be assigned to any group were categorized as unknown for each classification scheme.

### 2.3. Distance calculations

We rooted the maximum-likelihood tree for distance calculations by selecting samples that belong to bats and pangolin as an outgroup, namely EPI-ISL-412976, EPI-ISL-412977, and EPI-ISL-412860. We measured the distance from leaf to root for every leaf node that is present in the phylogenetic tree with the ete3 toolkit (Huerta-Cepas, et al. 2016).

### 2.4. Variant information processing

Mutations for each position relative to the reference genome (GenBank:NC_045512.2) were mapped catalogued in a table with a custom script. A table of all the mutations of selected sequences was created and ordered according to the phylogenetic tree of the corresponding genomes. Mutations that do not correspond to a nucleotide such as a gap or N were labeled as “Gap or N”; the other mutations were marked as Nongap. For variations that do not correspond to gap or N, respective nucleotides in the reference genome were obtained and added to the table. The GFF file of the reference genome (GCF_009858895.2) was extracted from NCBI Genome database. Open reading frame (ORF) information of each mutation was retrieved from the GFF file and added to the table. Positions that are not in the range of any ORF were labelled as “Non-coding region”. Codon information and position of each mutation in the reference genome were retrieved according to their respective ORF start positions and frame. In this process, reported frameshifts in ORF1ab (Dos Ramos, et al. 2004; Kelly and Dinman 2020) and the overlap between ORF7a 3’ and 7b 5’ ends were taken into account. Coding information was used to assign amino acid substitution information to the variations. Eventually, the variants were categorized as non-synonymous, synonymous, non-coding regions.

### 2.5. Migration analysis

The maximum-likelihood phylodynamic analysis was performed with Treetime (Sagulenko, et al. 2018) to estimate likely times of whole-genome sequences of SARS-CoV-2 by computing confidence intervals of node dates and reconstruct phylogenetic tree into the time-resolved tree. The slope of the root-to-tip regression was set to 0.0008 to avoid inaccurate inferences of substitution rates. With this model, we eliminated the variation of rapid changes in clock rates by integration along branches (standard deviation of the fixed clock rate estimate was set to 0.0004). The coalescent likelihood was performed with the Skyline (Strimmer and Pybus 2001) model to optimize branch lengths and dates of ancestral nodes and infer the evolutionary history of population size. The marginal maximum likelihood assignment was used to assign internal nodes to their most likely dates. Clock rates were filtered by removing tips that deviate more than four interquartile ranges from the root-to-tip versus time regression. JC69 model was used as General time-reversible (GTR) substitution models to calculate transition probability matrix, actual substitution rate matrix, and equilibrium frequencies of given attributes of sequences. The distribution of subleading migration states and entropies were recorded for each location through Augur trait module (sampling bias correction was set to 2.5). Closest child-parent pairs that do not go beyond their given locations were identified and evaluated as transmissions using Auspice (Hadfield, et al. 2018).

## 3. Results

### 3.1. Phylogenetic map of the virus subtypes

The first COVID-19 case in Turkey was reported on March 10^th,^ 2020, later than the reported first incidents in Asian and European countries. Since then, the number of cases increased dramatically. We used all the genomes available in the GISAID database as of May 1^st,^ 2020 and built a phylogenetic tree. After we filtered out the samples with incomplete date or location information, the total number of samples we eventually used was 15,277. The phylogenetic tree was built with the maximum likelihood method and a time-resolved tree was generated **(Figure 1)**. To verify the accuracy of the phylogenetic tree as well as to assess the distribution of well-characterized genomic features, we mapped several classification schemes on the tree; (i) S/L type (Tang, et al. 2020); (ii) D614G type (Bhattacharyya, et al. 2020); (iii) barcodes (Zhao, et al. 2020); (iv) six major clusters (Morais Júnior, et al. 2020). Although the methodologies of the clustering attempts were different between these studies, in general, the previously established groups were in line with our phylogenetic tree. Besides the already established clustering methods, we classified the clades based on the phylogenetic tree only. There are two levels of clustering; we termed phylo-clusters and phylo-subclusters. Small clusters were not taken into account (see Methods). The phylogenetic map of the virus genomes clearly shows the two major S and L type clades. As the ancestral clade, S-type is seen as limited in the number of genomes. 29 of the 30 isolates in Turkey were classified in the L-type group.

**Figure 1.**
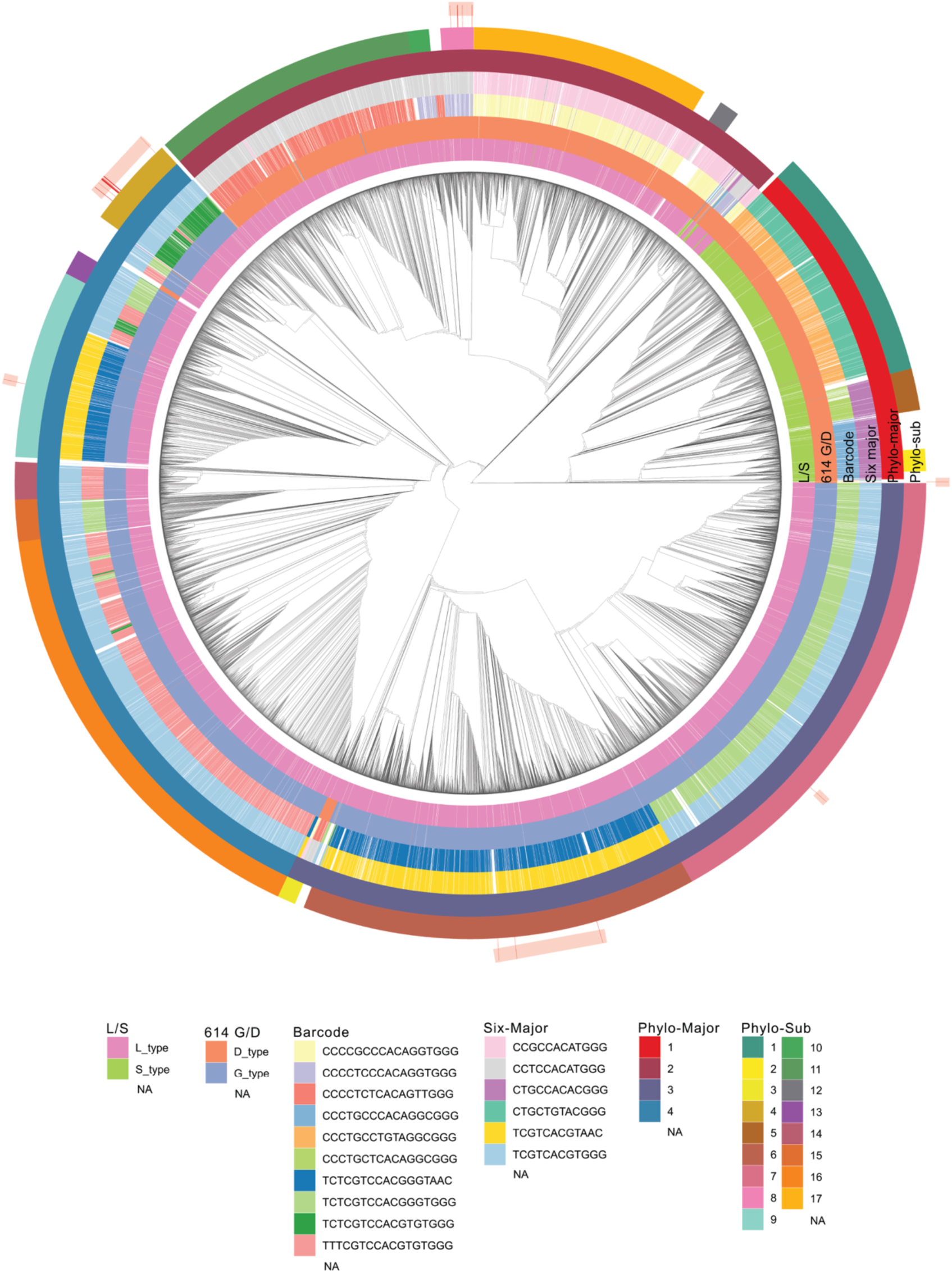
Phylogenetic tree of the 15,277 genomes retrieved from GISAID and their groupings. The time-resolved tree of SARS-CoV-2 appears in the center. Six clustering methods were used to assign 15277 sequences to the clusters. The clusters are represented as circular layers around the tree. The innermost shell (L/S) represents S and L type according to 8782th and 28144th positions in the nucleotide. 614 G/D represents the 614th amino acid of the Spike protein. Barcode shows the 10 major subtypes of seventeen positions in (nucleotide) multiple sequence alignment. Six-major clustering is based on 6 major subtypes of nucleotide combinations in particular positions. The fifth and sixth layers show Phylo-majors and sub-clusters, respectively. Samples obtained from Turkey are shown in the outermost shell and they are highlighted.

The samples from Turkey are dispersed throughout the phylogenetic tree **(Figure 1)**. The 30 samples are classified in 3 out of 4 different phylo-clusters and one remained unclassified. The dispersed groups suggested multiple independent introductions to the country. 7 of the 30 genomes encode aspartic acid (D) at the 614^th^ position of the Spike protein. The remaining 23 genomes encode glycine (G) in the same position. The D614G mutation is hypothesized to dominate because it enables smoother transmission of the virus (Bhattacharyya, et al. 2020). However, this correlation might simply be a founder effect which is basically the loss or gain of a genetic information when large population arise from a single individual.

### 3.2. A transient genome between S and L strain suggests early introduction

One of the genomes isolated in Turkey (EPI-ISL-428718) clustered together with the early subtypes of the virus. This isolate contains T at the position 8782, which is a characteristic of the S-type; however, it has T at the position 28144, which coincides with the L-type. Therefore, we characterized this sample as neither S-nor L-type. In the phylogenetic tree, this genome is placed between S and L strains, which suggests a transitioning genome from S to L strain **(Figure 2).** The number of variant nucleotides between this sample and root is lower relative to other Turkey samples. Phylogenetic placement in the earliest cluster, which is closer to the root, suggests that the lineage of EPI-ISL-428718 entered Turkey as one of the earliest genomes. By the time this sample was isolated in Turkey, the L-strain had started to spread in Europe, primarily in Italy. Although the isolation date of this early sample is one week later than the first reported case, the existence of an ancestral genome sequence suggests an earlier introduction of SARS-CoV-2 to Turkey.

**Figure 2.**
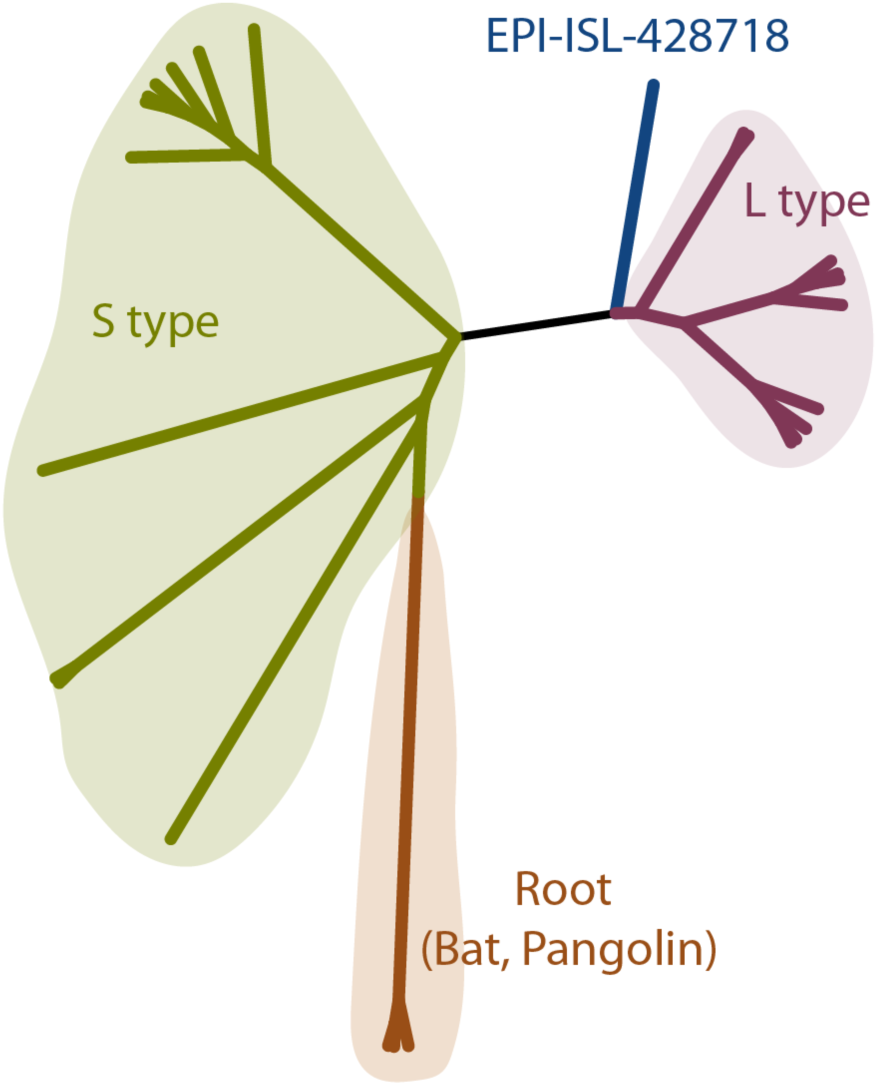
Phylogenetic tree of the transient type (EPI-ISL-428718) from S to L strain. The maximum likelihood tree was built with IQ-TREE. 10 S-type and 10 L-type sequences are randomly selected from the assigned samples. The tree was rooted at the genomes obtained from bat and pangolin.

### 3.3. Cluster profiles of the sample sets

Turkey has genome samples from at least three of the four major clusters. By taking the transitioning genome into account, samples of Turkey are genuinely scattered in the phylogenetic tree. Based on the groupings applied, we analyzed the relative abundances of the clusters in Turkey and other countries **(Figure 3A)**. The most samples of Turkey belong to cluster 4. Iran, Denmark and France are also enriched in cluster 4. Most European countries are enriched in cluster 3. Although Turkey has cluster 3 genomes, the fraction of them is lower compared to European countries. With the available genome sequences, the overall cluster profile of Turkey does not resemble any country. The divergence of the samples from to tree root was calculated for each sub-cluster. The sub-clusters observed in Turkey were analyzed along with the other countries **(Figure 3B)**. The divergence rates are comparable in general. However, within the same sub-clusters, virus genomes collected in Turkey have averagely more diverged than their relatives in other countries. The isolated genomes assigned to sub-cluster 4 and 8 show higher divergence rates in Turkey compared to the others in the same cluster (p-values: 0.00001 and 0.006, respectively, one tailed t-test between Turkey and the rest).

**Figure 3.**
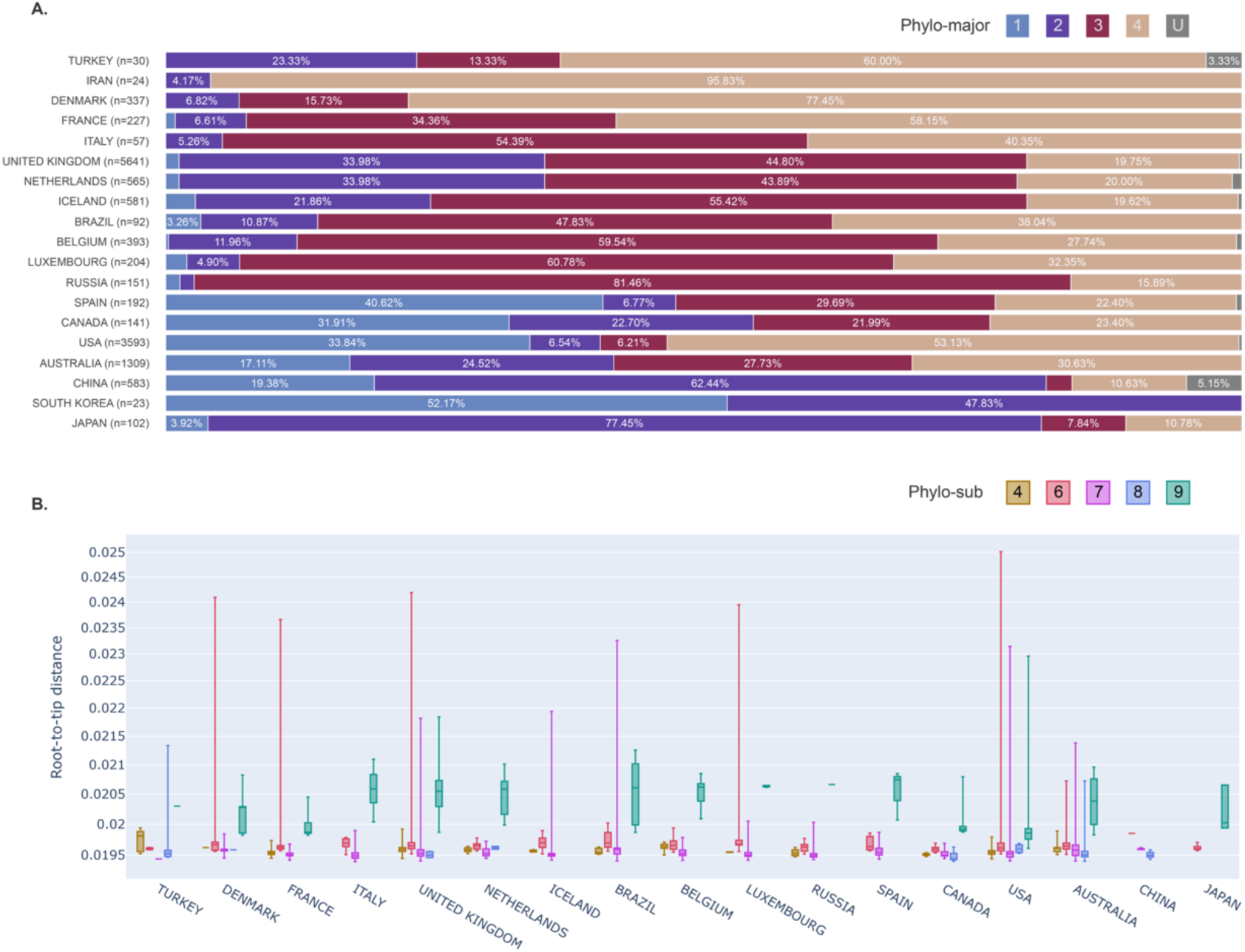
Phylo-cluster distribution and sub-cluster divergence. (A) Percentages of four major and unknown clusters across different countries. Unknown (U) samples are the ones that cannot be grouped with the generated clusters. (B) Root-to-tip distances of four phylo-sub clusters (4,6,7,8 and 9) found in Turkey, across different countries.

### 3.4. Mutation analysis of the genomes retrieved in Turkey

We used the Turkey isolates (30) to analyze their mutational patterns and their corresponding clusters. From the master tree, we pruned all the leaves except for the samples of interest. We rooted the subtree at the transitioning sample. We aligned the assigned clusters and all the mutations relative to the reference genome **(Figure 4)**, illustrating a correlation between the mutation pattern and the phylogenetic tree clades. Observation of no recurrence of a mutation shows that the multiple mutations are the results of founder effects.

**Figure 4.**
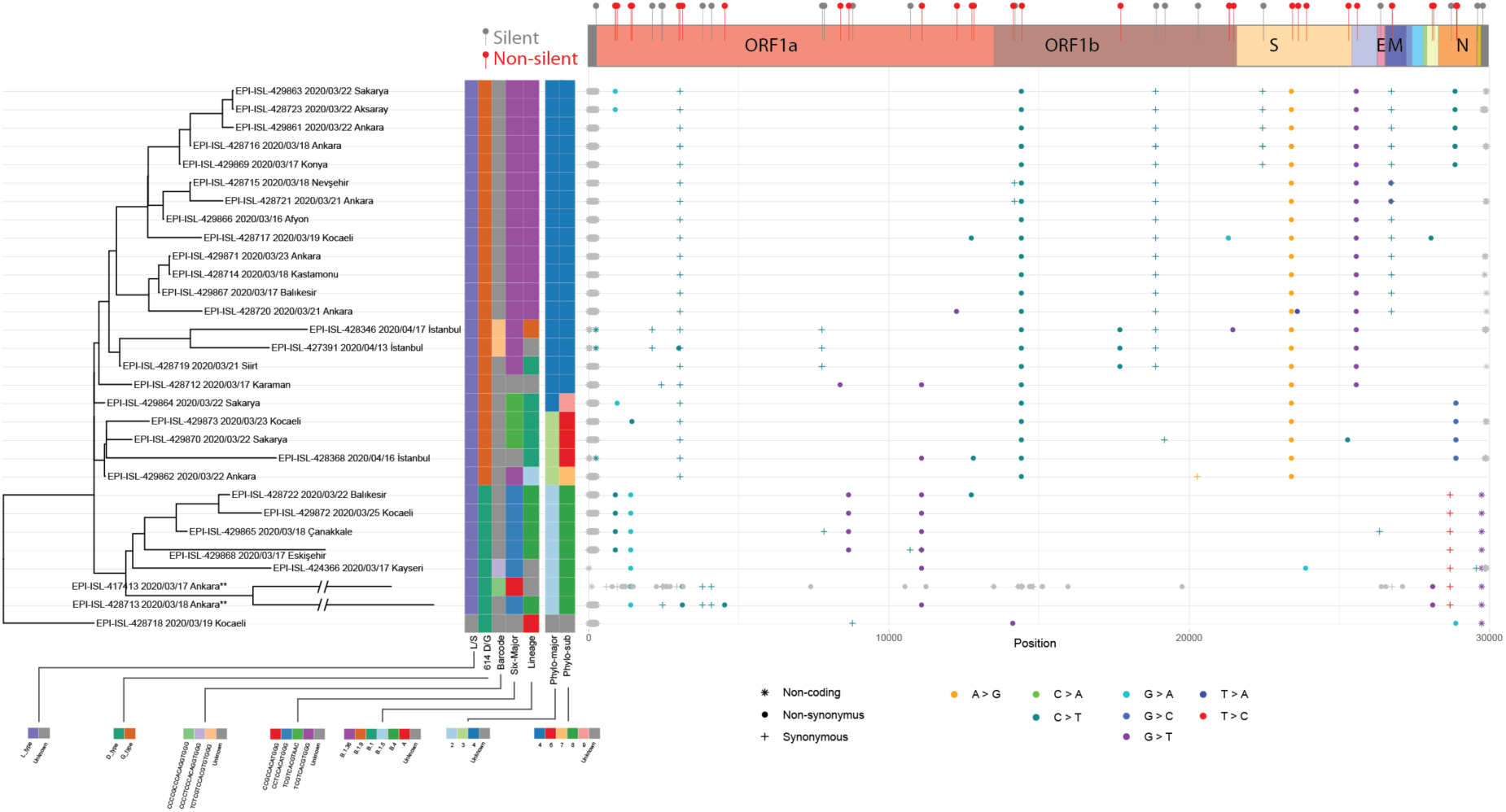
The mutation layout of the 30 samples from Turkey along with the phylogenetic tree and clusters. Phylogenetic tree (left) of SARS-CoV-2 samples sequenced in Turkey. Assigned subtypes of seven clustering methods are specified with different colors in the matrix. Dot-plot (right) of mutations detected in each genome aligned with the corresponding sample. Single nucleotide changes are colored and shaped based on the nucleotide change and synonymy. Gray color indicates that the mutation is either non-informative (ie, due to sequencing errors) or corresponds to a gap or an ambiguous nucleotide. Supplementary bar (top) provides the respective open reading frame information for mutations, and their effects on coding the amino acid. EPI-ISL-417413 had obvious sequencing errors, the mutations of this sampled were manually curated and non-informative ones were treated as ambigious mutations.

In total, 55 unique mutations were detected, 2 and 20 of which are non-coding and synonymous, respectively. Thirty-three unique amino acid substitutions are detected (Table 2). 23 out of 30 genomes we analyzed have the 614G mutation. The D614G mutation seems to have mutated with the two synonymous mutations in ORF1ab **(Figure 4)**. Besides 614G, three more amino acid substitutions were identified in the spike protein **(Table 2)**. G206A, T951I, G227S, S911F, A1420V, A3995F in ORF1a and V772I, T1238I in Spike protein, V66L in ORF5 and S54L in ORF8 were found to be specific to some isolates in Turkey **(Table 2)**. The most abundant amino acid substitutions (23/30) are P314L (ORF1b) and D614G (Spike), which are not specifically enriched in Turkey and dispersed worldwide. ORF1a V378I and ORF9 S194L are found in 7 and 6 of the 30 isolates, respectively, and show high frequency (15 folds with respect to general) in Turkey.

**Table 2.** Amino acid substitutions observed in 30 samples. The amino acid substitutions observed in Turkey are listed. The number of the overall substitutions were retrieved from CoV-GLUE database. The total number of genomes in the database was inferred from the D614G substitution which we found to be 63% of all the genomes. The substitutions that are observed at least in two isolates with enrichment factor greater than 2 are marked with *. (nt: nucleotide; aa: amino acid; EF: enrichment factor; sub: substitution)

| nt<br>pos | nt<br>sub | aa<br>pos | aa<br>sub | ORF | CoV-GLUE | Turkey<br>(30) | CoV-GLUE<br>fraction | Turkey<br>fraction | EF |  |
| --- | --- | --- | --- | --- | --- | --- | --- | --- | --- | --- |
| 881 | G > A | 206 | A>T | ORF1a | 2 | 2 | 0.00 | 0.07 | 565.60 | * |
| 884 | C > T | 207 | R>C | ORF1a | 52 | 4 | 0.00 | 0.13 | 43.51 | * |
| 944 | G > A | 227 | G>S | ORF1a | 1 | 1 | 0.00 | 0.03 | 565.60 |  |
| 1397 | G > A | 378 | V>I | ORF1a | 206 | 7 | 0.01 | 0.23 | 19.22 | * |
| 1437 | C > T | 391 | S>F | ORF1a | 27 | 1 | 0.00 | 0.03 | 20.95 |  |
| 2997 | C > T | 911 | S>F | ORF1a | 1 | 1 | 0.00 | 0.03 | 565.60 |  |
| 3117 | C > T | 951 | T>I | ORF1a | 1 | 2 | 0.00 | 0.07 | 1131.19 | * |
| 4524 | C > T | 1420 | A>V | ORF1a | 1 | 1 | 0.00 | 0.03 | 565.60 |  |
| 8371 | G > T | 2702 | Q>H | ORF1a | 22 | 1 | 0.00 | 0.03 | 25.71 |  |
| 8653 | G > T | 2796 | M>I | ORF1a | 55 | 4 | 0.00 | 0.13 | 41.13 | * |
| 11083 | G > T | 3606 | L>F | ORF1a | 2222 | 8 | 0.13 | 0.27 | 2.04 | * |
| 12248 | G > T | 3995 | A>S | ORF1a | 1 | 1 | 0.00 | 0.03 | 565.60 |  |
| 12741 | C > T | 4159 | T>I | ORF1a | 4 | 2 | 0.00 | 0.07 | 282.80 | * |
| 12809 | C > T | 4182 | L>F | ORF1a | 3606 | 1 | 0.21 | 0.03 | 0.16 |  |
| 14122 | G > T | 219 | G>C | ORF1b | 3 | 1 | 0.00 | 0.03 | 188.53 |  |
| 14408 | C > T | 314 | P>L | ORF1b | 10651 | 23 | 0.63 | 0.77 | 1.22 |  |
| 17690 | C > T | 1408 | S>L | ORF1b | 36 | 3 | 0.00 | 0.10 | 47.13 | * |
| 21304 | C > A | 2613 | R>N | ORF1b | 5 | 1 | 0.00 | 0.03 | 113.12 |  |
| 21305 | G > A | 2613 | R>N | ORF1b | 5 | 1 | 0.00 | 0.03 | 113.12 |  |
| 21452 | G > T | 2662 | G>V | ORF1b | 2662 | 1 | 0.16 | 0.03 | 0.21 |  |
| 23403 | A > G | 614 | D>G | ORF2 | 10691 | 23 | 0.63 | 0.77 | 1.22 |  |
| 23599 | T > A | 679 | N>K | ORF2 | 2 | 1 | 0.00 | 0.03 | 282.80 |  |
| 23876 | G > A | 772 | V>I | ORF2 | 1 | 1 | 0.00 | 0.03 | 565.60 |  |
| 25275 | C > T | 1238 | T>I | ORF2 | 1 | 1 | 0.00 | 0.03 | 565.60 |  |
| 25563 | G > T | 57 | Q>H | ORF3 | 4131 | 18 | 0.24 | 0.60 | 2.46 | * |
| 26718 | G > T | 66 | V>L | ORF5 | 2 | 2 | 0.00 | 0.07 | 565.60 | * |
| 28054 | C > T | 54 | S>L | ORF8 | 1 | 1 | 0.00 | 0.03 | 565.60 |  |
| 28109 | G > T | 72 | Q>H | ORF8 | 72 | 2 | 0.00 | 0.07 | 15.71 |  |
| 28854 | C > T | 194 | S>L | ORF9 | 220 | 6 | 0.01 | 0.20 | 15.43 | * |
| 28878 | G > A | 202 | S>N | ORF9 | 66 | 1 | 0.00 | 0.03 | 8.57 |  |
| 28881 | G > A | 203 | R>K | ORF9 | 3113 | 4 | 0.18 | 0.13 | 0.73 |  |
| 28882 | G > A | 203 | R>K | ORF9 | 3113 | 4 | 0.18 | 0.13 | 0.73 |  |
| 28883 | G > C | 204 | G>R | ORF9 | 3103 | 4 | 0.18 | 0.13 | 0.73 |  |

The mutational landscape represents the natural classifications of major and sub-clusters. These mutational footprints can be used to identify the clusters of the future genomes. The combinations of mutations can be used as barcodes to group upcoming virus genomes efficiently without a need for establishing evolutionary associations across lineages, which is a computationally expensive procedure considering the accumulating genomic data.

### 3.5. Trace of the spread

The number of mutations since December 2019 indicates that the SARS-CoV-2 genome mutates twice a month, on average. As genome sequencing reveals mutations, it enables a better understanding of the epidemiology by revealing the patterns of virus transmission. The time-resolved phylogenetic distributions of the genomes collected in Turkey suggested multiple independent sources of introduction **(Figure 5A)**. Out of the 30 genomes analyzed in this work, the earliest introduction seems to have originated from China. Other international imports include the US, Australia and Europe, probably from the UK. There is a connection between Saudi Arabia and the two cities in Turkey. Based on the model, this association is reciprocal. The Europe-based introductions are seen in the genomes isolated in Istanbul. Within Turkey, a transmission hub appears to be Ankara **(Figure 5B)**. The isolates in 5 cities are associated with genomes collected in Ankara **(Figure 5C).**

**Figure 5.**
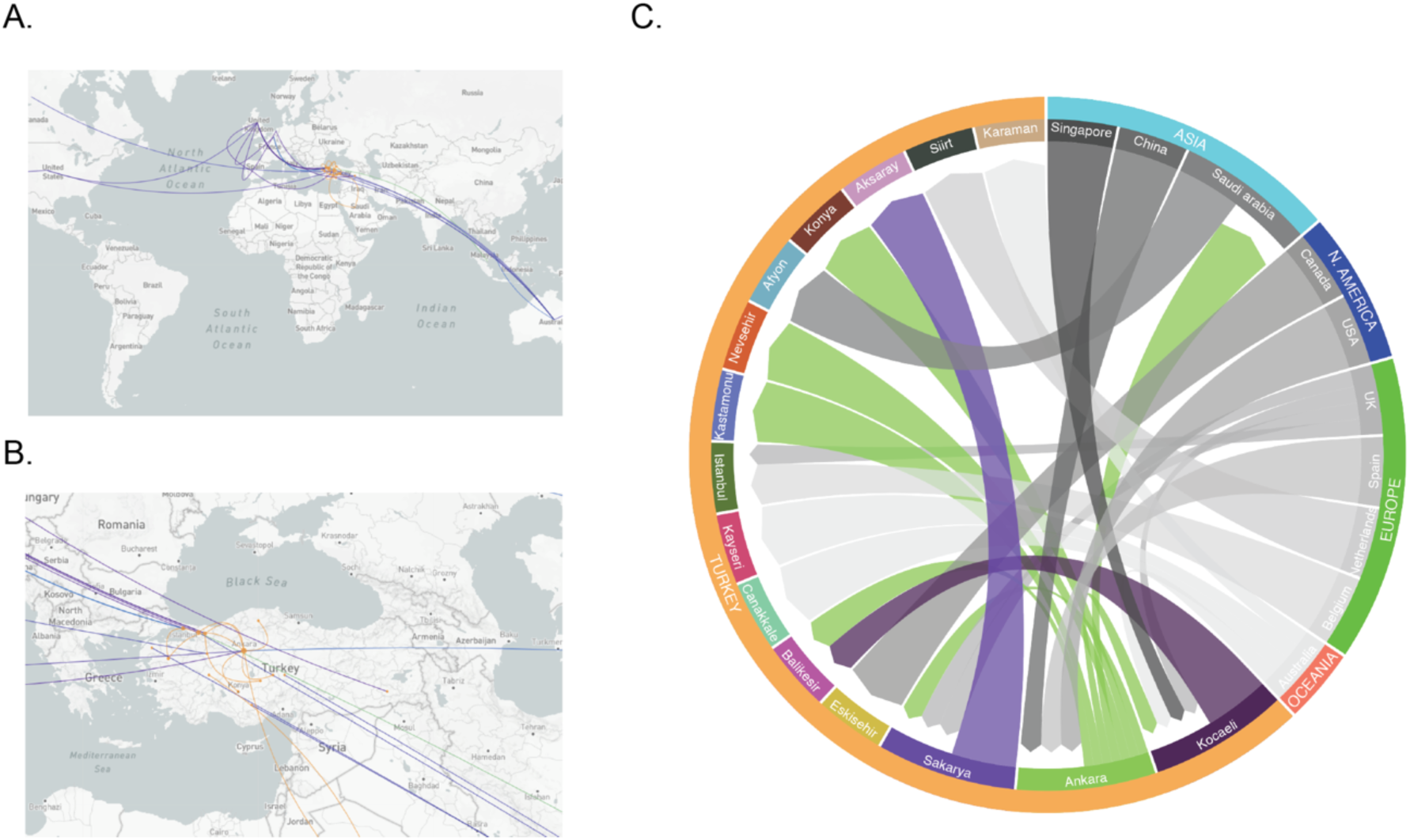
Epidemiological phylogenetic and transmission analysis of the isolates collected in Turkey. Sequences sampled between 2019-03-19 and 2020-04-24 were analyzed with Treetime and tracing between samples were visualized in Augur (version 6.4.3). (A) Closest (without internal nodes) leaves were used and assigned as transmissions were visualized on Leaflet world map using latitude & longitude information of locations. (B) Samples originated from Turkey were implied with orange points and connections while the network of samples originated from other countries demonstrated with blue lines and points. (C) Chord diagram was used as a graphical method to display inter-flow directed associations between origins and destinations of transmission data.

### 3.6. Web application to trace virus transmission

We have published a web application powered by Auspice (sarscov2.adebalilab.org/latest). We employed the front-end package (Auspice) that Nextstrain uses (Hadfield, et al. 2018). With increasing number of virus strains, not far from now, it will be infeasible to display the entire phylogenetic tree even in modern browsers. Nextstrain handles this problem by grouping the datasets based on the continents. As the aim of this platform is to trace the spread of virus genomes associated with Turkey, we will use representatives in the phylogenetic tree. The representative sequences will cover all the subtypes. The genomes of the samples collected in Turkey and their nearby sequences will be kept in the tree. With this approach, the web application will always contain the genome data from Turkey and necessary information of the subtypes with the representative sequences. An additional dimension we added to the application is that it enables to trace virus across the cities of Turkey. This approach is applicable to create a comprehensive platform for migration analysis for any country or region of choice.

## 4. Discussion

There are two most abundant lineages of isolates in Turkey: sub-clusters 4 and 8. If the 30 samples unbiasedly represent the overall distribution of the strains in Turkey, sub-clusters 4 and 8 might comprise approximately 80% of the genomes in the country. The high divergence of the samples in these sub-clusters in Turkey relative to their equivalents in other countries (Figure 3B) possibly suggests either or both of the two scenarios; (i) the viruses dominantly circulating in Turkey were introduced to the country later than other countries or (ii) this sub-cluster has been circulating in Turkey at a relatively higher rate than other countries and because of that, it is more likely to select the more diverged isolates by random sampling. Much more genomes should be sequenced and analyzed to gain more insight into virus evolution. It is essential to continuously follow up on the upcoming mutations when new samples are added to GISAID database.

The phylogenetic analysis of the circulating genomes in a country is necessary to identify the specific groups and their unique mutational patterns. The success of the COVID-19 diagnosis test kits, antibody tests and protein-targeting drugs possibly depend on genomic variations. For antibody tests, if a mutation affects protein recognition, the sensitivity of the test might drastically reduce. Therefore, mutation profiles of the isolates abundantly circulating in the country should be taken into account to modify these tests. As international travels are limited, viral genome profiles of the countries differ from each other, which is known as bottleneck effect. If international transmissions are kept being restricted, distinct cluster profiles might establish. Therefore, each country might need to develop their specific tests targeting the abundant genomes circulating in local.

We must note that sample distribution is not in line with the case distribution across Turkish cities. Due to this sampling bias as well as the low number of genomes, the spread history is undoubtedly incomplete. For instance, only 3 of the 30 samples were collected in Istanbul, which hosts approximately 60% of the COVID-19 cases. It is highly probable that Istanbul will be revealed as the central hub when additional genomes are sequenced. Moreover, there was no sample from Izmir, 3rd largest city. It should also be noted that the lack of a sufficient number of genomes could have resulted in indirect associations between the cities. More genomes are needed to complement this study with confidence.

The spread of the virus is traced by the personal declarations and travel history of the infected people. As SARS-CoV-2 genomes spread, they leave foot prints behind (mutations) allowing us to trace them. It is feasible to complement the conventional approach with genome sequencing in an unbiased way. Implemented feature of city-based tracing of the virus should be useful for authorities to take necessary measures to prevent spread. This approach will be automated with a standard pipeline. We aim to eliminate the technical limitations (because of the size) by applying filtering methods without losing any relevant information.

## Acknowledgments

This work, in part, is supported by the European Molecular Biology Organization (EMBO) Installation Grant (OA) funded by The Scientific and Technological Research Council of Turkey (TÜBİTAK). OA is additionally supported by International Fellowship for Outstanding Researchers Program, TÜBİTAK 2232 and BAGEP (Young Scientist Award by Science Academy, Turkey) 2019 grant. DÇ, ZK and BT are supported by the TÜBİTAK STAR program 2247-C.

We would like to thank all the healthcare workers who save lives during the COVID-19 pandemic. We thank the research groups who made the genome datasets available for accelerating research. So far, three groups in Turkey submitted genome sequences; 26 genomes were provided by the Ministry of Health (Fatma Bayrakdar, Ayşe Başak Altaş, Yasemin Coşgun, Gülay Korukluoğlu, Selçuk Kılıç); 3 submitted by the GLAB (Ilker Karacan, Tugba Kizilboga Akgun, Bugra Agaoglu, Gizem Alkurt, Jale Yildiz, Betsi Köse, Elifnaz Çelik, Mehtap Aydın, Levent Doganay, Gizem Dinler); 1 submitted by Erciyes University (Shaikh Terkis Islam Pavel, Hazel Yetiskin, Gunsu Aydin, Can Holyavkin, Muhammet Ali Uygut, Zehra B Dursun, İlhami Celik, Alper Iseri, Aykut Ozdarendeli).

We thank Dr. Barış Süzek and Dr. Batu Erman for their helpful comments on the manuscript. We would like to acknowledge Cem Azgari for his contributions throughout the project. We thank Molecular Biology Association for their leadership in taking the initiative of forming a pool of volunteers in COVID-19 testing. Finally, we would like to thank the members of Ecology and Evolutionary Biology Association in Turkey for fruitful discussions regarding the preliminary analysis of the SARS-CoV-2 genomes.

## Authors’ contribution

OA conceived the study, designed the analysis, interpreted the results and wrote the first draft. AB generated the multiple sequence alignments, Bİ generated the visualization pipeline with auspice. BS generated the clusters based on the phylogenetic tree and plotted cluster graphs. DÇ, ZK and BT assigned previously identified clusters to the genomes, visualized the clusters aligned with the tree and identified mutations per sample. All authors contributed to manuscript writing and revising.

